# Multi-omics data integration via novel interpretable k-hop graph attention network for signaling network inference

**DOI:** 10.1101/2022.09.16.508281

**Authors:** Ruoying Yuan, Jiarui Feng, Heming Zhang, Yixin Chen, Philip Payne, Fuhai Li

## Abstract

With the advent of sequencing technology, large-scale multi-omics data have been generated to understand the diversity and heterogeneity of genetic targets and associated complex signaling pathways at multiple levels in diseases, which are critical targets to guide the development of personalized precision medicine. However, it remains a challenging task to computationally mine a few essential targets and pathways from a large number of variables characterized by the multi-level multi-omics data. In this study, we proposed a novel interpretable k-hop graph attention network model, k-hop GAT, to integrate the multi-omics data to infer the essential targets and related signaling networks. We evaluated the proposed model using the multi-omics data, i.e., genetic mutation, copy number variation, methylation, gene expression data, of 332 cancer lines; and the experimentally identified essential targets. The validation and comparison results indicated that the proposed model outperformed the GAT and graph convolutional network (GCN) models.

## Introduction

With the advance of sequencing technology, multi-omics data have been generated to understand the genetic heterogeneity and complex signaling pathways at multiple levels in diseases. Compared with solo-omics data analysis, the integration of the multi-omics datasets can 1) provide a holistic view of complex and multi-level biological processes, 2) increase the statistical power to identify the molecular mechanism which involves causal and vital molecular targets and signaling pathways. For example, the multi-omics datasets were generated in the cancer genome atlas (TCGA)^1,2,3^ project to characterize the molecular diversity and heterogeneity of cancer patients. In addition, the Cancer Cell Line Encyclopedia (CCLE)^4^ project has generated multi-omics data of about 1000 different cancer cell lines, which can indicate the different and distinct dysfunctional signaling targets and pathways. The Genomics of Drug Sensitivity in Cancer (GDSC) project also associated the multi-omics data of cancer cell lines with some drug sensitivity^5^. Also, the multi-omics data were generated in the Religious Orders Study and Memory and Aging Project (ROSMAP)^6,7^ project to study the genetic variants and dysfunctional signaling pathways in Alzheimer*’*s disease (AD). The multi-omics data, i.e., genetic mutations and copy number variations (genetic), methylations (epigenetic), gene expression (RNA-seq), proteomics (protein), measure and characterize the multi-level molecular genotype of diseases. These multi-omics datasets have been useful to identify essential disease biomarkers and uncover some dysfunctional signaling pathways. However, it remains a challenging task to integrate the multi-omics data and mine the vital signaling targets and signaling pathways, which can guide the personalized disease management and the development of personalized precision medicine or cocktail medicines, from a holistic view.

On the other hand, to guide the analysis of the large-scale multi-omics datasets, experimental screening using RNAi or crispr technology has also been conducted to identify the essential targets that can affect the phenotype of diseases, like cancer cell proliferation and migration. For example, large-scale target screening has been conducted to experimentally identify essential signaling targets to inhibit the tumor cell growth in the DepMap^8^ project. The valuable multi-omics datasets in CCLE and genetic screens, i.e., genetic effects on the phenotypes, in DepMap provide the basis that can catalyze the development of novel computational models to facilitate the development of precision medicine.

The multi-omic data analysis remains a challenging task^9^. A comprehensive review of existing multi-omics data integration analysis models was reported^9^. Specifically, these models were clustered into a few categories, like similarity, correlation, Bayesian, multivariate, fusion and network-based models^9^. Among the network-based models, the paradigm^10^ model is one of the most widely used models, which was built based on the factor graph. Compared with the traditional computational models, in the recent study^11^, the DeepDep model was developed based on the combination of deep belief network (DBN)^12^ and auto-encoders^13,14^, to integrate the multi-omics data of CCLE and TCGA for the DepMap target effect prediction. Though it achieved good prediction accuracy, the model interpretation is still limited to the perturbation analysis, which cannot interpret and uncover the subsequent dysfunctional signaling pathways of the essential molecular targets.

It is well known that proteins within cells work coordinately as a systematic network and modules to regulate a set of complex biological processes and dysfunctional signaling pathways in complex diseases, like cancer^2^. For example, 10 core signaling pathways were systematically using the multi-omics data of TCGA, which indicated the diverse and essential targets of these signaling pathways for different cancer subtypes. Moreover, a set of signaling pathways, like KEGG^15,16^, wikiPathways^17^, and protein-protein interaction (PPI) database, like BioGRID^18,19^, STRING^20,21^, have been reported and publicly accessible. Thus, graph neural network (GNN)-based models^22–24^ can naturally represent the signaling flow/interactions on the signaling networks; and the latent status of individual proteins will be affected by their multi-omics data (features) and also the interacting proteins (neighbors) on the signaling network (graph)^25,26^. More importantly, using the attention mechanism it is possible identify the essential targets and subsequent signaling pathways. Thus, in this study, we proposed to integrate the multi-omics data using a novel k-hop graph attention network (k-hop GAT); and to infer the essential targets and subsequent signaling networks. The detailed introduction of datasets, model architecture and evaluation were presented in the following sections.

## Materials and Methodology

### Multi-omics datasets of cancer cell lines

The multi-omics data of cancer cell lines were collected from Cancer Cell Line Encyclopedia (CCLE)^4^ project (see **Table 1** and **Table 2**). Four types of data, i.e., gene level transcriptomics, methylation, copy number variation, genetic mutations, were used as the features of proteins nodes in the deep graph neural network (GNN) models.

**Table 1.** Datasets used in this study.

| Dataset/Database | Description | URL link |
| --- | --- | --- |
| CCLE | Multi-omics data of ~1,000 cancer cell lines (including gene level <u>transcriptomics, methylation, copy number variation, genetic mutations</u>), which will be used to estimate the activity of individual proteins. (<u>Used as features of protein nodes on the KEGG signaling graph in the GNN models</u>). | https://sites.broadinstitute.org/ccle/datasets |
| DepMap | <u>Gene effect scores, which are derived from CRISPR knockout screen for each cancer line. The negative scores indicate cell growth inhibition effects of the gene knockout. (Used as the label to train the GNN models).</u><br><u>Meta-data of DepMap cancer cell lines. (Used to align the DepMap cancer cell lines with CCLE).</u> | https://depmap.org/port al/download/ |
| KEGG | The KEGG signaling pathways will be merged to construct the signaling network. (Used as the graph (with 1985 protein nodes) in the GNN models). | https://www.genome.jp/kegg/ |

**Table 2.** Statistics of CCLE multi-omics data.

| CCLE multi-omics data | # of genes/proteins | # of cancer cell lines |
| --- | --- | --- |
| Gene expression | 19177 | 1393 |
| Mutation | 48270 | 1032 |
| Copy number | 20671 | 980 |
| methylation | 21338 | 846 |
|  | 1985 (proteins on the KEGG signaling pathways) | 332 (# of cancer cell lines with all 4 types of omics data.) |

### KEGG signaling network

In the KEGG database, there are 311 signaling pathways, extracted using the graphite R package^27,28,15^. These signaling pathways were merged to construct the signaling network. Then a signaling network with 1,985 proteins, with all the 4 types of data, was used as the graph (to calculate the adjacency matrix) in the GNN models (see **Table 1**). The Fisher*’*s exact test^29,30^ was used for the signaling pathway enrichment analysis.

### DepMap gene effect data

In the DepMap data^8^ (see **Table 2**), the gene effect score (scores indicate cell growth inhibition effects of the gene knockout) was used to indicate the cancer cell growth inhibition effects of specific genes. In this exploratory study, we selected 100 targets with the lowest DepMap priority scores (inhibiting tumor cell growth, as the positive targets) (the priority score of each gene is defined for cancer types, higher score means the gene is more important in the given cancer type (not cell line level)); and selected 50 targets with highest DepMap scores (promoting tumor growth, as the negative targets); and randomly selected 50 targets. Finally, we will model the gene effect scores of the 200 genes on the 332 cell lines, as the label to train the GNN models.

### Model architecture of the K-hop Graph Attention Network (GAT) model

The attention mechanism was introduced to graph neural network in graph attention network (GAT)^31^. One limitation of the GAT model is the lack of expressive power because it is bounded by the Weisfeiler-Lehman(1-WL) test^32^. Instead of using 1-hop neighbor, we proposed to improve the GAT model by using k-hop neighbors that can aggregate information from genes k-steps away on the signaling network/graph. Specifically, to identify the k-hop neighbors, the shortest path distance was used in our model. The k-hop model can be more efficient to distinguish nodes with different nodes in the k-step neighborhood/field (see **Fig. 1**).

**Figure 1.**
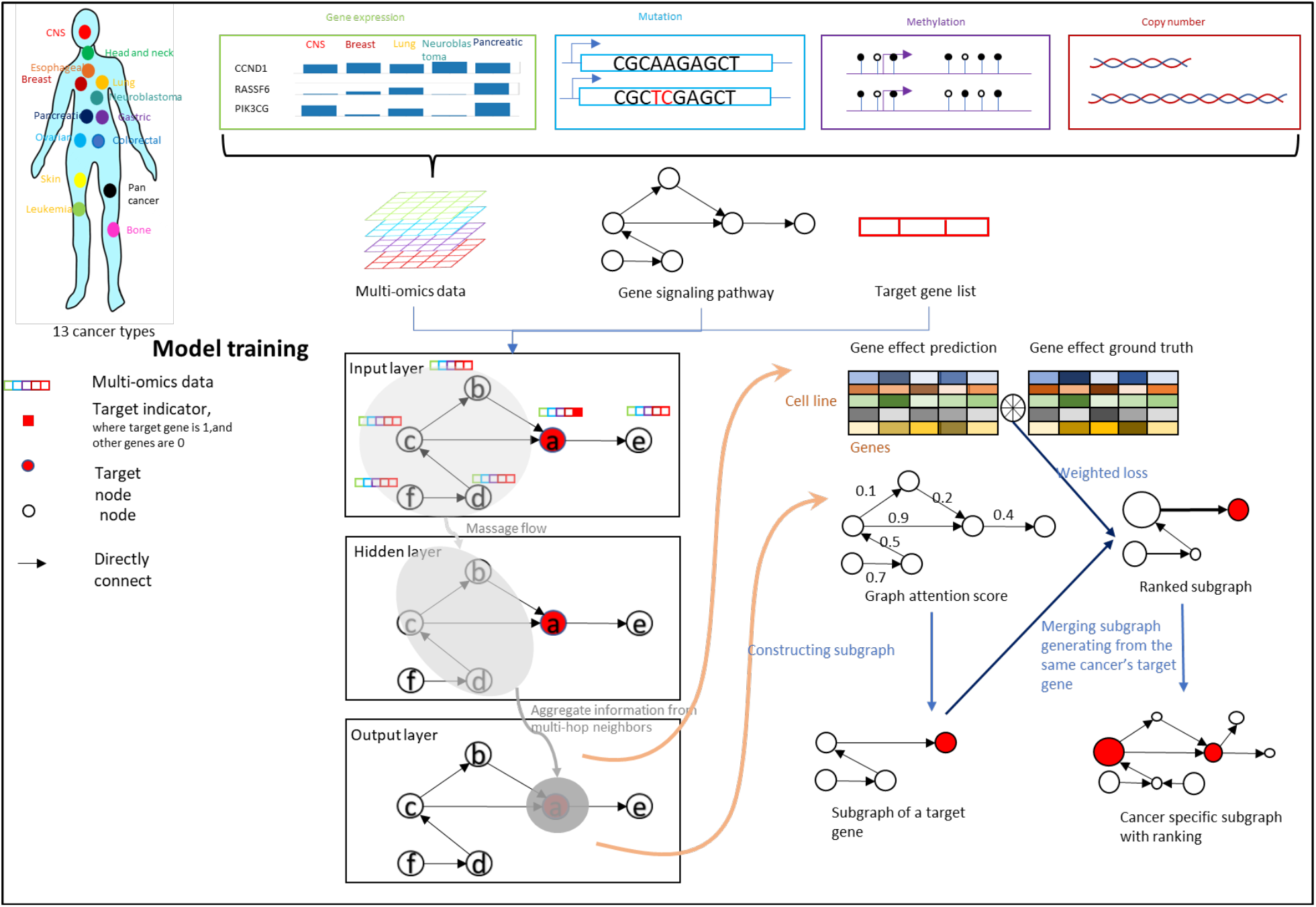
Overview, architecture and signaling-flow of the proposed k-hop GAT model.

As seen in **Fig. 1**, the model input variables are: *X* ∈ *R*^*n*×*d*×*5*^, which represents the multi-omics data of *n* cell lines and *d* genes. For each gene in a cancer cell line, there are 4 types of genomics data (i.e., gene level transcriptomics, methylation, copy number variation, genetic mutations); and the 5^th^ element is used to indicate if it is a target to be modeled (to predict its gene effect score). *A* ∈ *R*^*d*×*d*^ is the adjacent matrix of the selected KEGG signaling network. To make the model interpretable and avoid potential noisy effects, the receptive field pooling for a given target was added. In this study, the proposed model used the attention scores of 1 and 2 hop neighbors, and has 2 layers. Thus, the receptive filed is 4-hop neighbors. In another word, the 4-hop neighbors can pass signaling to the given target. Specifically, we implemented the receptive field pooling by adding a mask to output before pooling, as defined in the following equations:

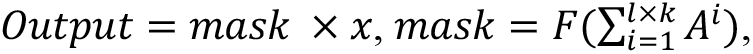

where K is the number of k-hops, and l is number of layers of the model. F is a function that set all the non-zero value to 1. based on shortest path distance kernel is the set of nodes that have the shortest path distance from node v less than or equal to K.

We denoted 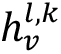as the output representation of all the k-th hop neighbors of node v at layer l, 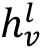as the output representation of node v at layer l. Then, the K-hop signaling flow (or message passing) is defined as follows:

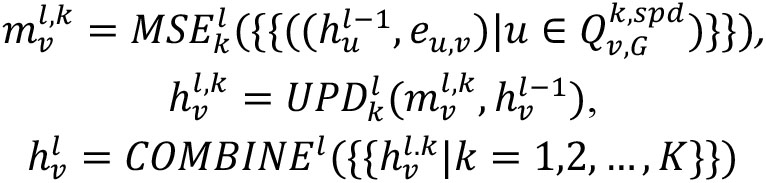

where 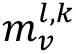is the message from the k-th hop neighbors of node v to node v at layer l at, 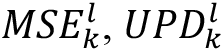 are message and update functions at hop k at layer l (see **Fig. 2**).

**Figure 2.**
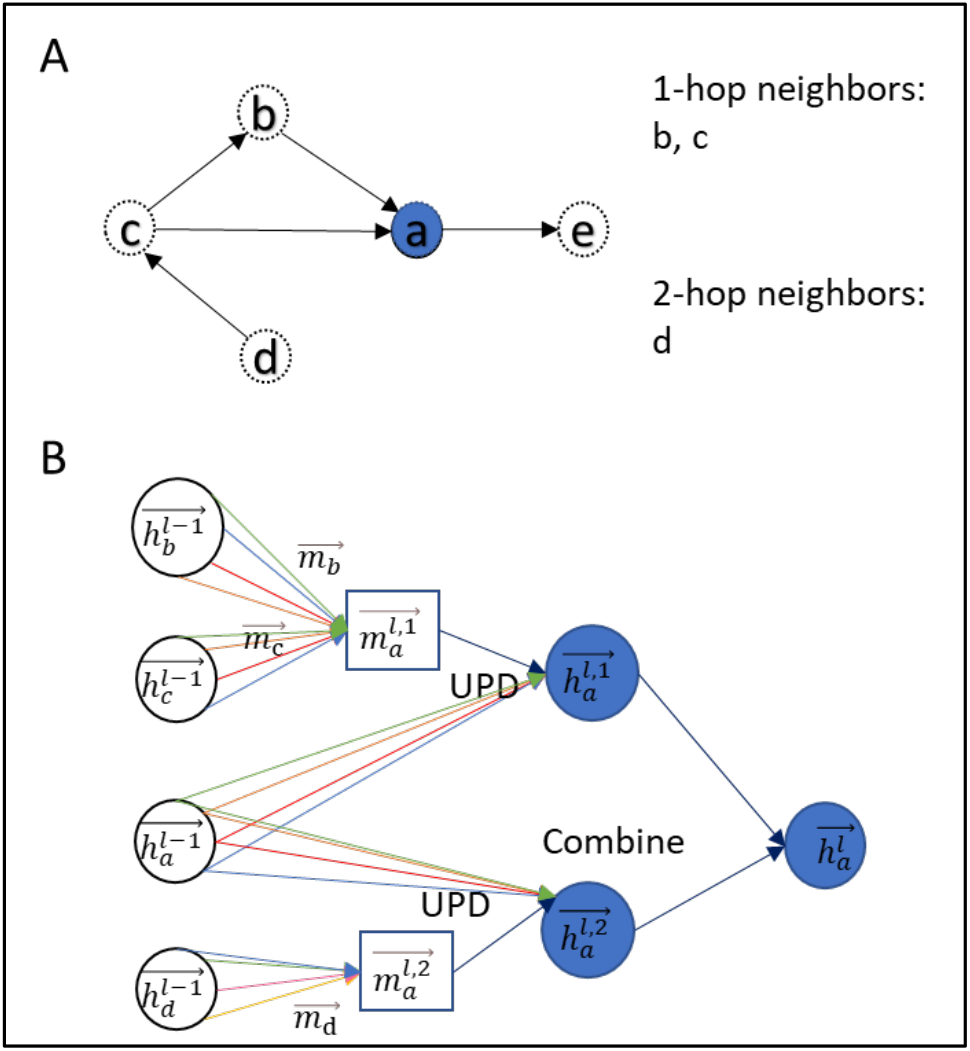
Illustration of the signaling-flow on the k-hop signaling neighbors. (A): For a node v in graph G, the 1- and 2-hop neighbors 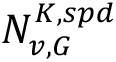 of v based on shortest path distance. (B) The 2-hop message passing flows for the graph (A), different color implies different heads. The color of edges indicates the different heads.

### Essential signaling network inference/subnetwork extraction

After training the model, the attention scores on the graph for each cell line will be obtained, which will be used to infer the essential subsequent signaling flows related to the target genes. In total, there are 332 (cell lines) x 200 (target genes) x 4 (heads) attention scores. For each head in each cancer cell line for a given target, we first choose the one-hop and two-hop neighbors with attention scores higher than a given threshold. For the two-hop neighbors, since there might have more than one path, the signaling path with the largest product of the edge attention scores will be selected, because larger product of attention represents more signaling flows on this path. Considering that we used 2 layer 2-hop GAT model, thus the receptive field is 4 hop neighbors. Then, we use the similar method to find the 3 and 4 hop neighbors of the target starting from the 1-2 hop neighbors. The selected paths from multiple heads will be merged to construct the signaling network of given targets on specifical cancer cell lines. Moreover, for each given cancer type, the selected signaling pathways for a set of given targets and a set of cancer cell lines (belonging to the same cancer type) will be merged as the cancer type specific dependence signaling network.

### Scoring target importance

It is an important task to score the potential importance of targets on the selected signaling networks. In this study, we defined the target importance score, for each cancer cell line, using the average attention scores of multiple heads. For a given cancer type, the target importance score is defined as the weighted average of attention scores across all the related cancer cell lines, where the weight for each cancer cell line is defined as:

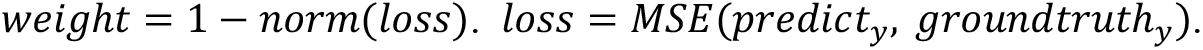

## Results

### Model hyperparameters

The model was implemented using Pytorch and geometric. The Adam optimizer was used. The initial learning rate = 0.0005 with a minimum of 1e-6 to terminate the training. We empirically set the training epochs = 40. We set the K=2 and l=2 empirically because 3-hop will have millions of edges, which is computationally expensive and will cover almost all the nodes on the whole signaling network. The hidden layer size is 128; the leakyRelu parameter was set to 0.2; the output dimension is 100; and reduced to 1 dimension after the receptive field pooling. The 5-fold cross-validation was used. The mean square error (MSE) and correlation (between the predicted gene effect scores and the experimental gene effect scores) were used as the loss function.

### Model performance and comparison

**Table 3** shows the MSE and correlation values on the training and testing dataset. As seen, the model generated similar performance on the training and testing data respectively. We further compared the proposed model with the other two widely used models, i.e., GAT and GCN (see **Table 4**). As seen, the proposed model outperformed the GAT and GCN models significantly.

**Table 3.** MSE and correlation values of the proposed model on the 5-fold cross-validation datasets.

| Number of folds | MSE on Training data | MSE on Testing data | Correlation on Training data | Correlation on Testing data |
| --- | --- | --- | --- | --- |
| 1<sup>st</sup> fold | 0.1387 | 0.1345 | 0.7090 | 0.7047 |
| 2<sup>nd</sup> fold | 0.1269 | 0.1272 | 0.7241 | 0.7224 |
| 3<sup>rd</sup> fold | 0.1260 | 0.1307 | 0.7246 | 0.7228 |
| 4<sup>th</sup> fold | 0.1458 | 0.1474 | 0.6874 | 0.6960 |
| 5<sup>th</sup> fold | 0.1435 | 0.1450 | 0.7005 | 0.6951 |
| Mean | 0.1361 | 0.1366 | 0.7094 | 0.7088 |

**Table 4.** Model comparison with graph attention network (GAT) and graph convolutional network (GCN).

| Models | Average MSE on training data | Average MSE on Testing data | Average correlation on Training data | Average correlation on Testing data |
| --- | --- | --- | --- | --- |
| The proposed model | 0.1361 | 0.1366 | 0.7094 | 0.7088 |
| GAT | 0.1740 | 0.1765 | 0.5523 | 0.5506 |
| GCN | 0.1613 | 0.1607 | 0.6411 | 0.6431 |

### Inferred subsequent signaling networks of selected essential targets in breast, lung and pancreatic cancers

**Fig. 3** shows the inferred subsequent signaling networks of the selected essential targets in breast, lung and pancreatic cancers. We just empirically selected the breast, lung and pancreatic cancer types as examples. As seen, the three cancer types have diverse and different essential signaling networks. There are 119, 133 and 116 targets selected in the breast, lung and pancreatic cancer respectively. As seen in the venn-diagram in **Fig. 4**, there are many targets are specific to the cancer types. Moreover, we conducted the literature search about the identified essential signaling targets. Interestingly, many of the top-ranked targets have been associated with supportive literatures, which indicated the importance of the proposed model. Due to the page limit, the related references were not cited here (and attached as a supplementary file). For example, literatures have been reported the important roles of *PIK3CG, RASFF6, MYB, SPP1, THRA, ZAP70, NFKB2, CDKN1A, AKT2, TFDP1, DNM1L, CCNE1, CDK1, RCAN1, FGFR1, CSF1, DVL3, RAC1, CDKN2A, RBCK1* in breast cancer. For the lung cancer, the *MYB, CDKN2A, CDK4, TFDP1, DVL3, TFAM, PIK3CG, E2F4, NFKB2, CDK1, AKT2, SPP1, RCAN1, SLC2A2, RAB8A, KLF2, CCNE1, HHIP, STK11, XIAP, EGR3* were reported. For the pancreatic cancer, the *RASSF6, PKM, MYB, CDC25A, CCNB1, FOSL1, E2F2, CDKN2A, FOXO1, CCND3, RBCK1, NFKB2, RBL1, MAGI2, LDHA* were reported. The results indicated that the proposed model can identify additional essential targets using the anchor targets and the integration of the multi-omics data and signaling pathways.

**Figure 3.**
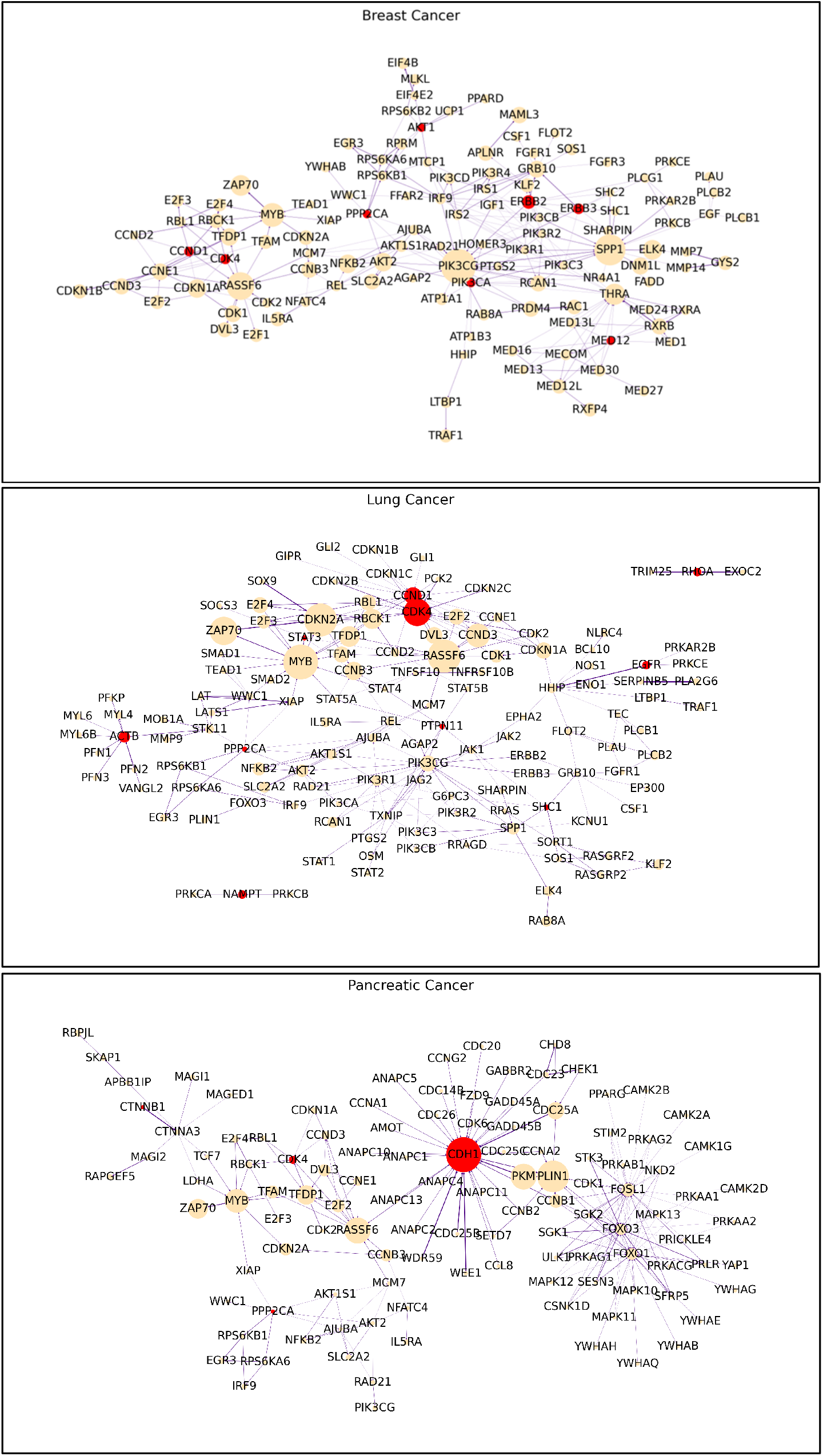
Inferred subsequent signaling networks of given selected essential targets (red) in breast, lung and pancreatic cancers. The node size and edge size are proportional to target importance scores and edge attention scores respectively.

**Figure 4.**
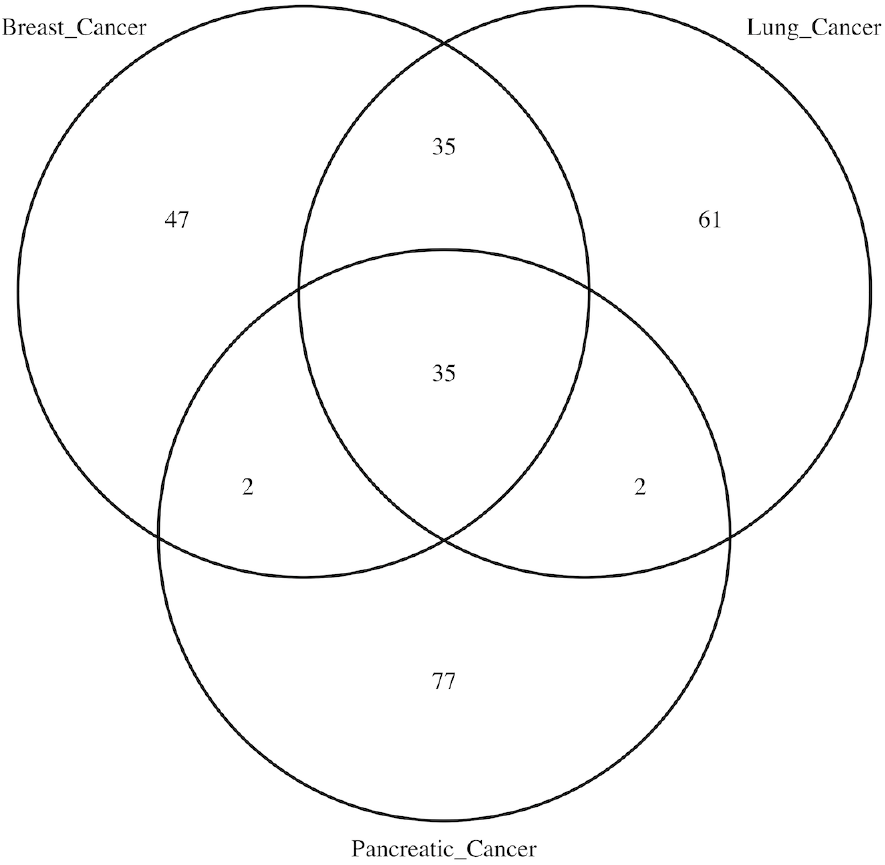
Common and unique targets on the inferred essential signaling networks of breast, lung and pancreatic cancer types.

## Conclusion

Large-scale multi-omics datasets have been generated for studying the genetic diversity and dysfunctional signaling targets and subsequent signaling pathways, which is the critical basis for the development of personalized precision medicine. However, the multi-omics data integration and interpretation remain challenging tasks. Novel computational models are needed urgently. In this study, we developed a novel k-hop graph attention network (k-hop GAT) model to resolve these challenges in multi-omics data integration and interpretation. The evaluation of the proposed model using the multi-omics data 332 cancer lines and the experimentally validated targets indicated that the proposed model outperformed the GAT and GCN models. Moreover, we defined the target importance scores and subsequent signaling network inference using the attention scores on the background signaling graph. Thus, we conclude that the K-hop GAT model is efficient to integrate and interpret the multi-omics data for inferring essential signaling networks.

## Limitations

There are some limitations of this study. First, we only selected a subset of the targets validated in DepMap to train and evaluate the proposed model. In the following work, we will train the model using all these targets to uncover the potential holistic view of the essential signaling pathways for each individual cancer cell lines or cancer subtypes. The essential signaling targets and pathways will be important for identifying novel medicines or drug cocktails, inhibiting multiple vital targets on the signaling network, as novel treatment regimens of specific cancer subtypes. Moreover, it is critical to make the model more interpretable by identifying the causal factors, i.e., which level of genetic dysfunction triggered the activation of the targets and their subsequent signaling pathways. It can further narrow down the search space of the potential therapeutic targets toward personalized precision medicine. The third limitation is that the analysis was conducted using the known KEGG signaling pathways. Thus, it is interesting and necessary to identify novel causal factors by using large-scale protein-protein interaction networks. In future work, we will improve the proposed models to resolve these challenges to make the multi-omics data well integrated and more interpretable.

## Acknowledgment

This work is partially supported by the Children*’*s Discovery Institute (CDI) M-II-2019-802 to Li.

